# Genetic Adaptation in New York City Rats

**DOI:** 10.1101/2020.02.07.938969

**Authors:** Arbel Harpak, Nandita Garud, Noah A. Rosenberg, Dmitri A. Petrov, Matthew Combs, Pleuni S. Pennings, Jason Munshi-South

**Affiliations:** Department of Biological Sciences, Columbia University, New York, NY; Department of Ecology and Evolutionary Biology, University of California, Los Angeles, CA; Department of Biology, Stanford University, Stanford, CA; Department of Biological Sciences, Fordham University, Armonk, NY; Department of Ecology, Evolution and Environmental Biology, Columbia University, New York, NY; Department of Biology, San Francisco State University, San Francisco, CA

## Abstract

Brown rats (*Rattus norvegicus*) thrive in urban environments by navigating the anthropocentric environment and taking advantage of human resources and by-products. From the human perspective, rats are a chronic problem that causes billions of dollars in damage to agriculture, health and infrastructure. Did genetic adaptation play a role in the spread of rats in cities? To approach this question, we collected whole-genome sequences from 29 brown rats from New York City (NYC) and scanned for genetic signatures of adaptation. We tested for (i) high-frequency, extended haplotypes that could indicate selective sweeps and (ii) loci of extreme genetic differentiation between the NYC sample and a sample from the presumed ancestral range of brown rats in northeast China. We found candidate selective sweeps near or inside genes associated with metabolism, diet, the nervous system and locomotory behavior. Patterns of differentiation between NYC and Chinese rats at putative sweep loci suggests that many sweeps began after the split from the ancestral population. Together, our results suggest several hypotheses on adaptation in rats living in close proximity to humans.

## Background

Urbanization has the potential to drive dramatic ecological and evolutionary consequences for wildlife [1, 2]. The most commonly examined evolutionary responses to urbanization are changes in gene flow and in the intensity of genetic drift [3, 4, 5, 6]. A small but growing number of studies have also examined adaptive evolution in urban environments, including morphological adaptations to urban infrastructure [7, 8], adaptive life history changes, thermal tolerance to warming cities [9, 10], and behavioral changes [11]. The recent adoption of genomic scans to identify loci under positive selection can help generate hypotheses about adaptive phenotypes in cities [12, 13].

Commensal rodents—particularly house mice (*Mus musculus*), black rats (*Rattus rattus*), and brown rats (*Rattus norvegicus*)—are the most widespread urban mammals besides humans and a notorious threat to urban quality of life [14, 15]. Recent analyses have revealed some of the relationships between invasive urban rodent populations that spread around the world with humans [16, 17, 18, 19] and the influence of heterogeneous urban environments on gene flow [20, 21]. Much less is known about the role of natural selection in the success of commensal rodents in cities.

Brown rats have considerably less genetic diversity compared to house mice, possibly due to a population bottleneck ∼20,000 years ago [22]. More recently, they have experienced major range expansions, presumably due to their association with agrarian, and later urban, human societies [19]. After reaching Europe around 500 years ago, brown rats rapidly became a prominent urban pest and then spread throughout Africa, the Americas, and Australia as a side effect of European colonialism in the 18th and 19th centuries. These introductions to coastal cities, followed by rapid industrialization and urbanization, potentially exerted strong selection on rat populations. Urban environments changed most dramatically from the late 19th into the 20th century–a period that spans around 500 rat generations. A recent example is found in evidence for a significant change in rat cranial shape in New York City over a 120 year period [23].

It is likely that selection influenced a number of traits in these expanding populations. For example, multiple rat populations exhibit resistance to first-generation anticoagulant rodenticides (such as Warfarin) associated with nonsynonymous substitutions in the VKORC1 gene [24], though resistance to recent second-generation anticoagulants is less widespread and may not be monogenic [25]. Comparisons of Asian and European rats have suggested that many immune-response genes are highly differentiated between these regions, potentially due to selection related to disease pressures [26]. Rats in NYC evolved longer noses—which have been interpreted as adaptations to cold, and shorter upper tooth rows—which were interpreted as adaptations to higher quality, softer diets [23]. Although no study to date has examined the genomic signatures of selection in urban rats, studies of another rodent in NYC, the white-footed mouse (*Peromyscus leucopus*), provide additional hypotheses on the traits subject to selection in urban rodents. Namely, immune response, detoxification of exogenous compounds, spermatogenesis, and metabolism genes were over-represented among putatively-selected regions [27, 28, 29]. This same *Peromyscus* population exhibited shorter toothrows consistent with a change in diet [30]. These findings suggest that some genetic adaptations in urban rodents have arisen in response to increased disease pressure in dense urban settings, increased exposure to pollutants and shifts to novel diets.

Here, we perform a genomic scan for adaptation in an urban rat population. We search for signatures of recent selective sweeps—long haplotypes at a high frequency [31, 32, 33]— in a sample of rats from NYC. We also search for adaptive changes by identifying regions of extreme differentiation between our NYC sample and a second sample from northeast China—the presumed ancestral range of the species [16, 19]. We find evidence for recent selective sweeps near genes associated with metabolism of endogenous compounds, diet, apoptotic processes, animal organ morphogenesis and locomotory behavior.

## Results

To look for selective sweeps in NYC brown rats, we sequenced whole genomes of 29 rats trapped throughout Manhattan, New York City, USA. The animals were chosen to represent the geographic distribution of Manhattan rats while excluding genetic relatives (**Fig. 1**) [20]. The mean coverage was ≥ 15x for each of the 29 rats. The mean nucleotide diversity was 0.168 ± 0.003% (point estimate ± standard error), slightly lower than the 0.188 ± 0.0001% estimated for a population at the presumed ancestral range of brown rats in Harbin, China (**Supplementary Materials**) [22].

**Figure 1:**
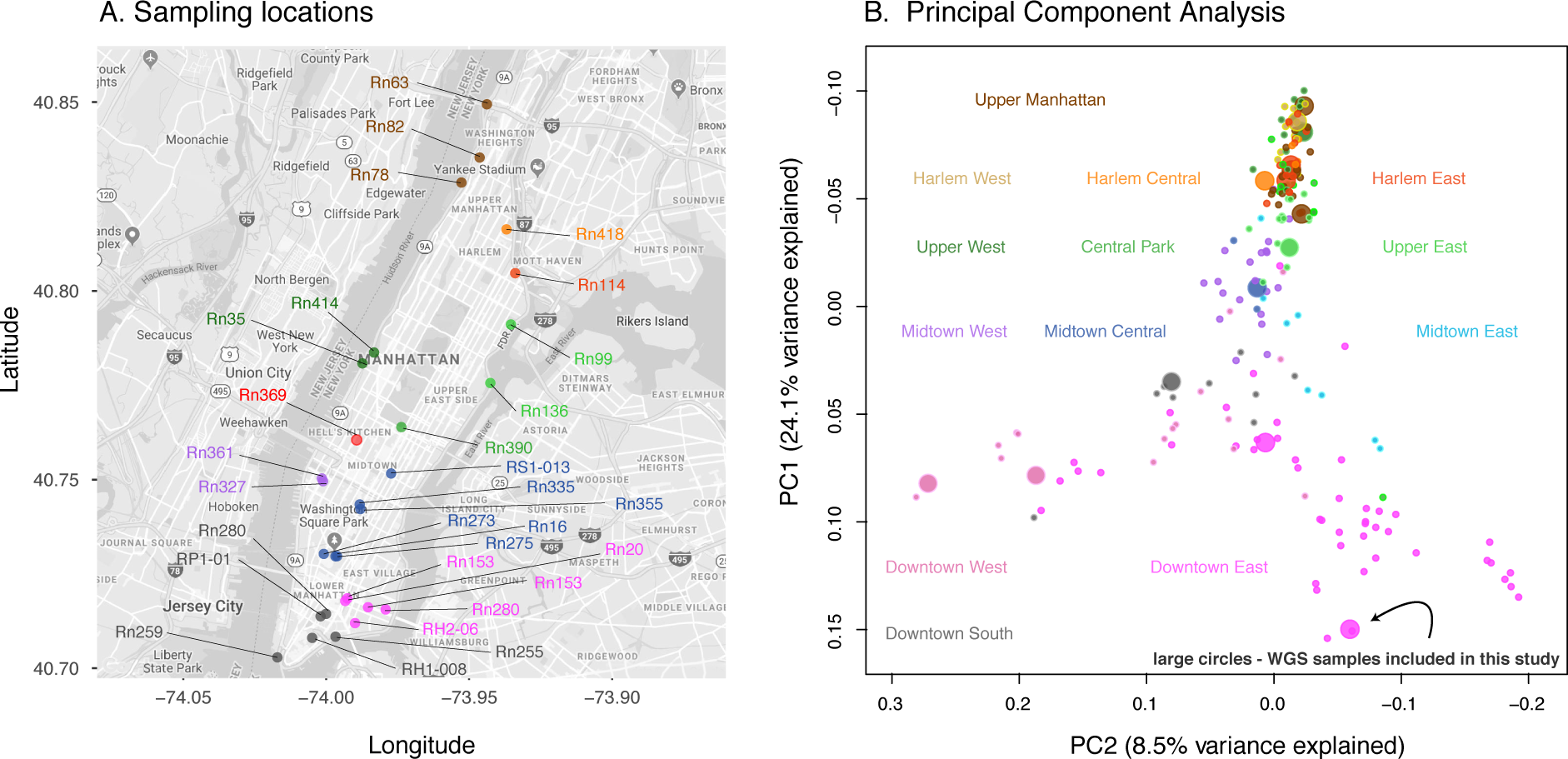
Sampling locations and genetic population structure of NYC brown rats. **(A)** Sampling sites of the 29 rat whole-genome samples used in our selection scans. Color coding corresponds to NYC neighborhoods as in panel (B). **(B)** Principal component analysis of 198 rats reanalyzed from Combs et al. [20], with the 15 rats included in both the Combs et al. [20] study and this study represented by large circles. In both panels, color coding corresponds to NYC neighborhoods.

We searched for selective sweeps using the methods G12 [34, 35] and H-scan [36]. G12 and H-scan both measure the homogeneity of haplotypes in the sample around a focal SNP (**Methods**).

High haplotype homogeneity is a signature of a recent (or sometimes even ongoing) selective sweep. We chose to use G12 and H-scan because they do not require prior phasing of the genotype data—a potential challenge in a small sample. Instead of relying on an assumption of perfectly phased data, G12 and H-scan measure *multi-site genotype homogeneity*. It has been recently argued that—under a wide range of selection parameters and demographic histories—methods using unphased data are almost as statistically powerful as methods based on perfectly phased data [35, 37]. G12 and H-scan were correlated (Spearman *ρ* = 0.52; *p <* 2.2 × 10^*-*16^), and they tended to peak at similar regions of the genome—suggesting that they detect similar signals of frequent, long segregating haplotypes (**Fig. 2; Fig. S1A**).

**Figure 2:**
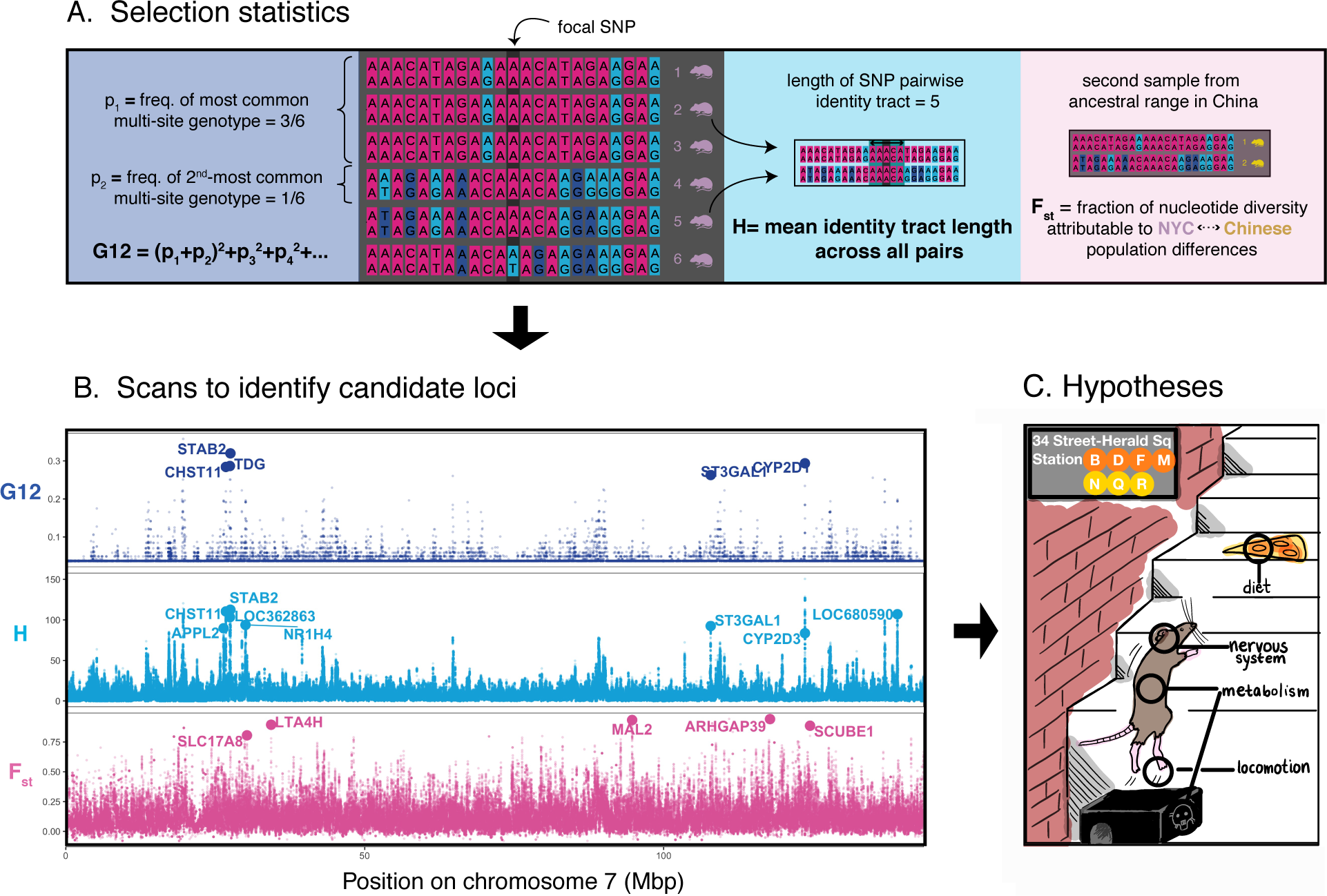
Scanning for signals of adaptation. **(A)** Three selection statistics were used to identify signals of selection. Two of the statistics, G12 and H-scan, detect loci with low haplotype diversity (high homogeneity)—a signature of recent or ongoing selective sweeps. G12 is defined similarly to the homozygosity of multi-site genotypes (in a window of a fixed number of SNPs)—a sum of squared frequencies of multi-site genotypes—except that the two most frequent genotypes are grouped (summed and squared) together. In the illustrated example, the most common multi-site genotype is shared by rats 1,2 and 3. The second-most common genotype is only carried by one rat. H is the mean length of the maximal pairwise identity tract containing the focal SNP—where the mean is taken across all pairs in the sample. In the illustrated example, the tract is 5 SNPs-long for rats 2 and 5. Finally, *F*_*st*_ was used to measure genetic differentiation between the NYC sample and a sample from the presumed ancestral range of brown rats in northeast China. Extreme differentiation at a locus may also be indicative of selection since the populations’ split. **(B)** Values of the three statistics along chromosome 7. G12 and H-scan peak at similar regions of the genome, suggesting that both statistics pick up on similar signals of frequent, long segregating haplo-types. Large circles denote the locations of candidates: top-scoring loci that are also less than 20kb away from protein-coding genes (names of the corresponding genes are noted). Corresponding plots for all other autosomes can be found in Supplementary File S1. **(C)** Functions associated with candidate genes suggest traits that may have been subject to selection in NYC rats, including diet, the nervous system, metabolism and locomotion.

A common approach to evaluating significance of evidence for selection is to test if an empirical value of a statistic is a likely sample from a specified null distribution. The null distribution can be estimated using simulations of neutral evolution under an inferred demographic model. We initially took this approach, but found that the simulated null distribution of G12 was far from the empirical one, suggesting that the null model was poorly calibrated (**Fig. S2**; **Supplementary Materials**).

We therefore took an alternative approach of focusing on the most extreme empirical values of the selection statistics as putative targets of adaptation (**Fig. S7**; we note that we still report the neutral simulation results (**Table S1**) and p-values computed using the simulation-based null distribution (**Table S6**)). We calculated G12 and H-scan values across the whole genome (**Files S3,S4**). We identified the 100 top-scoring loci with each method, iteratively masking regions around previously-called loci to avoid correlated signals due to linkage disequilibrium (**Supplementary Materials**). Multi-site genotypes are visibly more homogeneous in top-scoring loci (**Fig. 3B,C**) than in random loci (**Fig. 3A**). In some of the top-scoring loci, we observe a single common haplotype—consistent with the expectation under a hard selective sweep [38, 39, 40, 41] (**Fig. 3B**). In many others, more than two common haplotypes are segregating in the population—consistent with the expectation under a soft selective sweep [42, 43] (**Fig. 3C; File S2**).

**Figure 3:**
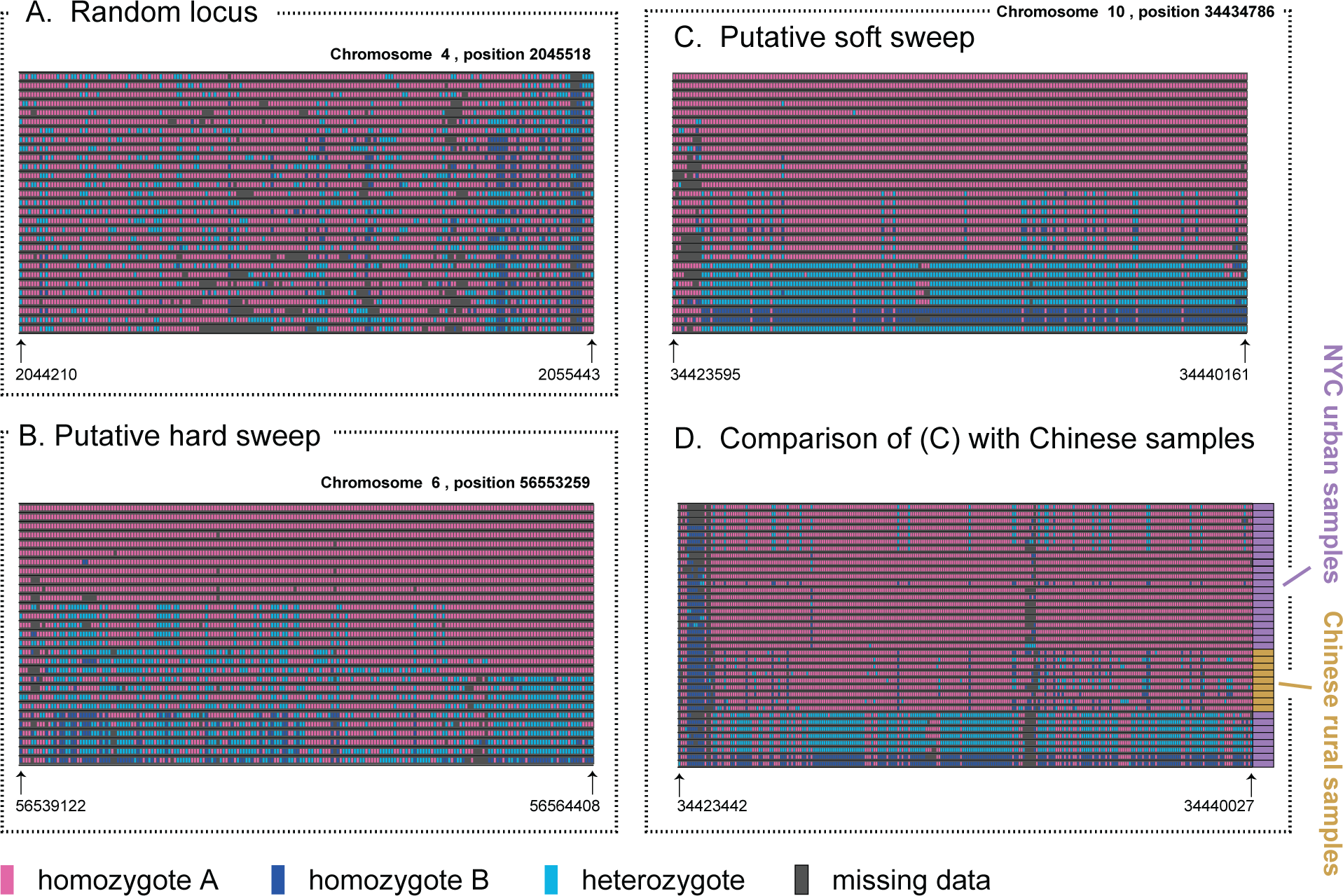
Multi-site genotype structure. Each row confers to a multi-site genotype of one rat. **(A)** At a randomly-chosen locus, there is no clear clustering of multi-site genotypes. **(B)** An example of a locus that was detected as a putative target of selection with H-scan, in which there is one multi-site genotype frequent (11/29) in the NYC sample. This pattern is consistent with a hard selective sweep. **(C)** A candidate locus identified with G12 in which there are more than two frequent multi-site genotypes—consistent with a soft selective sweep. **(D)** A comparison of the NYC samples to a sample of 9 rats from the presumed ancestral range of the species in China, around the same focal site as in panel (C). Multi-site genotypes are relatively heterogeneous in the Chinese sample where homogeneity is observed in the NYC sample. This observation is consistent with a recent onset of selection after the split from the ancestral population in China.

To gauge the timing of putative selective sweeps, we compared the multi-site genotypes of our sample to those of 9 rats from Harbin, Heilongjiang Province, China—the presumed ancestral range of the species [22]. The Chinese genotypes are often heterogeneous at the loci most homogeneous in the NYC sample (**Fig. 3D**). Only 5/17 of the H-scan candidates show significantly elevated H-scores in the Chinese sample (**Fig. S4; File S6**). These patterns are consistent with most of these putative selective sweeps occurring in ancestors of the NYC sample after the split from the ancestral population in China (Multi-site genotype visualizations for all G12 and H-scan candidate loci can be found in **File S2**).

We next took a partially complementary approach to find candidate targets of adaptation: searching for regions of extreme genetic differentiation between our NYC sample and the Chinese sample. We measured differentiation using mean per-SNP *F*_*st*_ [44] across SNPs in 10kb sliding windows (**File S5**), and again focused on the 100 top-scoring loci genome-wide (**Table S3**). In the **Supplementary Materials** we discuss the relationship between top-scoring loci of one statistic and scores in the other. In short, if a genotype homogeneity peak is due to a recent selective sweep that has occurred after the split from the ancestral population, then we expect high *F*_*st*_ values in the same region. Indeed, *F*_*st*_ values are slightly elevated (80th percentile for H-scan, 76th percentile for G12) in genotype homogeneity candidates (**Fig. S1A**). This pattern is consistent with a selective sweep elevating local differentiation from the Chinese population, although we note that even if no selective sweeps have occurred, then *F*_*st*_ is expected to be somewhat elevated from the genome-wide baseline at loci ascertained for high homogeneity in the NYC sample [45].

We next asked if we could associate biological functions to these top-scoring loci. For each candidate locus, we identified a single closest gene. The top-scoring G12 and H-scan loci are farther from protein-coding genes than is expected under a permutation-based null (Wilcoxon *p <* 0.004 for both, but not for *F*_*st*_; **Fig. 2C**; **Supplementary Materials**). This result may suggest that genic sweeps are rare—at least among loci identified with our methods.

Moving forward, we only considered protein-coding genes ≤ 20*kb* away from top-scoring loci as potential targets of adaptation—leaving 19, 17 and 32 candidates for G12, H-scan and *F*_*st*_ respectively (**Tables S1-S3**). One G12 candidate lies close to a a cluster of Cytochrome P450 genes (It is specifically closest to CYP2D1, but there are five CYP2D genes ≤ 50*kb* away). CYP-encoded enzymes help metabolize endogenous compounds and detoxify exogenous chemicals [46] and have been associated with Warfarin resistance [47, 48]. This gene family has expanded substantially in rodents [49].

Several of the genes—3 of 17 for H-scan and 2 of 19 G12 candidates—are olfactory receptor genes. Changes in the olfactory nervous system can alter odor perception and behaviors driven by precise chemical cues—and have been suggested to be targets of adaptation in numerous mammalian populations [50]. However, this is not a significant enrichment compared with a null generated by permuting the location of the candidate loci along the genome (Fisher’s Exact Test; **Supplementary Materials**).

We do, however, find Gene Ontology (GO) biological processes [51] that are enriched among top-scoring genes. Among the top fourteen G12 values inside genes (coresponding to 0.27 expected false discoveries, **Methods**), three biological processes are associated with at least three different genes: apoptotic process, animal organ morphogenesis and axon guidance (**Fig. 4**; largely overlapping results using H-scan instead of G12 are shown in **Fig. S3**).

**Figure 4:**
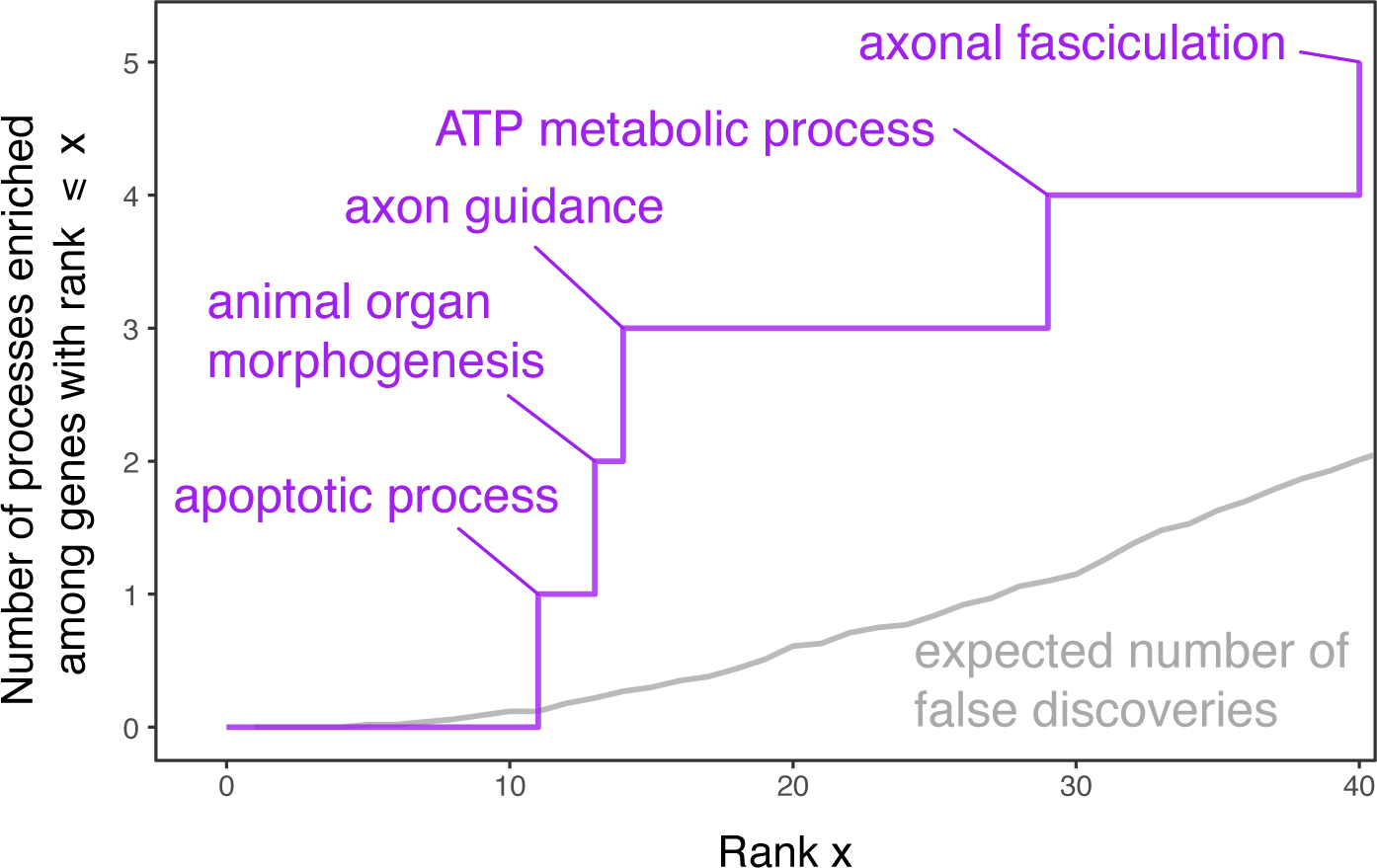
Biological processes enriched among 40 top-scoring genes. This figure focuses on candidate targets of adaptation arising from a comparison of G12 values in genes. Each gene was given the maximum score of all focal SNPs in the gene, and genes were ranked by this maximal score. We consider Gene Ontology (GO) biological processes associated with at least three top-scoring genes as enriched, and the y-axis shows their cumulative count. Blue text labels show GO biological processes associated with exactly three of the (x value) top-scoring genes. The gray line shows the expected number of false (biological process) discoveries among the top X candidates, estimated based on permutations of gene scores.

One potential candidate that we had considered before performing our scans was VKORC1, as previous research has pointed to selective sweeps at this locus in rodents following the broad application of Warfarin as a rodenticide [24, 52]. The Chinese and NYC rats were highly differentiated at a 10kb window centered on VKORC1 (98th chromosomal percentile of *F*_*st*_). At the same time, multi-site genotypes around VKORC1 did not appear homogeneous, and G12 and H-scan are close to the chromosomal medians (**Fig. S5**). Zooming in on the VKORC1 coding sequence, we found that no known resistance-conferring alleles appear to have fixed in the population (**Supplementary Material**), and, further, that the data contained no nonsynonymous SNPs. To our knowledge, all of the experimentally-confirmed resistance-conferring variants in VKORC1 are nonsynonymous [24]. We detected 10 intronic SNPs, and two synonymous SNPs (in codon 68 coding for a Histidine and in codon 82 coding for Isoleucine)—both previously observed in NYC rats [53] (**Fig. S6**).

## Discussion

We performed scans for adaptation in NYC rats and identified the top-scoring loci as potential targets of recent selective sweeps. Near or within top-scoring loci, we find genes associated with the nervous system, metabolism apoptotic processes and morphogenic traits.

One trait of central interest as a target of adaptation is rodenticide resistance. Some of the genes we identified near candidate loci (e.g. the CYP2D gene cluster and AHR; **Tables S1-S5**) may be associated with rodenticide resistance. In VKORC1—which we had a priori considered as a possible target of such adaptation—we find no evidence for a recent selective sweep (though levels of differentiation between Chinese and NYC rats in the genomic region surrounding VKORC1 were high, see **Fig. S5**). We also detect no nonsynonymous polymorphisms in the gene (**Fig. S6**; but see [53] for evidence of two nonsynonymous variants associated with resistance segregating at low frequencies in NYC). Warfarin became the rodenticide of choice in New York in the late 1950s [54], but was gradually phased out in the late 1970s and replaced by more effective second-generation anticoagulant rodenticides (SGAR) after high levels of resistance were reported from Europe [55, 56] and the United States [57], and specifically New York [58]. It is possible that sweeps that have occurred during these few decades are too old to be detectable with our methods, although resistance-associated VKORC1 polymorphisms are still segregating in some European populations [24]. Another possibility is that a VKORC1 sweep did not occur in the NYC population.

Natural populations of brown rats have been reported to show resistance to SGAR as well [59, 60]. VKORC1 may play a role in SGAR resistance—but likely a smaller role than it does for first-generation anticoagulants [25]: lab rats homozygous for the resistance mutation Y139C showed lower mortality in response to some SGARs (difenacoum and bromadiolone), but not others (brodifacoum, difethialone and flocoumafen) [61]. At the same time, rats resistant to the SGAR bromadialone showed increased expression of a suite of CYP genes [62]. We may therefore hypothesize that the signals of adaptation we observe near the CYP2D gene cluster could underlie changes in response to SGAR through the clearance of exogenous compounds. However, to our knowledge, levels of anticoagulant (including warfarin) resistance in NYC rats have not been quantified in recent decades.

These genes could also be under selective pressure from other environmental toxins, inflammatory responses, or novel dietary items. For example, one of our candidate genes, AHR, has been implicated in adaptation to polychlorinated biphenyl (PCB) contamination in killifish and tomcod in the Hudson and other rivers in the area [63, 64]. On a possibly related note, our GO analysis identified the nervous system genes as a possible target (**Fig. 4,S3**). This putative adaptation may also relate to neuronal or hormonal responses to the environment, but may also underlie behavioral changes.

Other candidate genes and biological processes highlighted in **Figs. 4,S3 and Tables S1-3** are associated with many phenotypes in humans and lab rodents that are also plausible targets of selection. However, the sheer number of phenotypic associations for each gene makes it difficult to generate clear hypotheses about phenotypes under selection in wild urban rats. In addition, in the absence of a sample from rural counterparts of our urban study population, we cannot discern whether any of the adaptations are driven by the urban environment with certainty. Even with this caveat, there are some potentially promising leads that follow from this study. As one example, the gene CACNA1C (7th highest score among genes for both G12 and H-scan, **Tables S4-5**) has been repeatedly associated with psychiatric disorders in humans, and the effects may be modulated by early-life stress [65]. Thus, rat behavioral phenotypes associated with anxiety (such as anti-predator responses or responses to novel stimuli) are promising areas for future investigation. CACNA1C has also been demonstrated to influence social behavior and communication in rats [66]. CACNA1C and multiple other top-scoring genes like FGF12, EPHA3 and CHAT (**Table S4**) may also affect locomotion in rodents [67, 68]. The gait or other locomotory phenotypes could have also undergone adaptive changes, given that urban rats must move through a highly artificial, constructed environment that differs markedly from naturally vegetated habitats.

Given that urban rats are so closely associated with human city dwellers, future work might explicitly address whether rats respond to the pressures of city living that are also experienced by humans. One striking commonality between urban humans and rats is their diet. Today, the human urban diet contains an increasingly large proportion of highly processed sugars and fats that lead to a number of public health concerns. Some of these health concerns could conceivably apply to rats as well [69, 70, 71]. Indeed, several of the candidate genes we identified, ST6GALNAC1 (G12, **Table S1**), CHST11 and GBTG1 (H-scan, **Table S2**), are linked with oligosaccharide or carbohydrate metabolic processes.

The main approach we used to identify targets of adaptation was to search for long haplotypes segregating at a high frequency. The results of our scans should be viewed only as preliminary hypotheses for future testing: in the absence of a sensible null model, the specificity of our genotype-homogeneity outlier approach remains unclear. In addition, the power of our approach to detect genetic adaptation as a whole is limited; it is most powerful for ongoing or recent selective sweeps—and the definition of recency here is also a function of the sample size (as larger samples can reveal more recent coalescence events) [35, 72]. Other modes of adaptation, such as polygenic adaptation [73, 74, 75, 76], may be common in NYC rats, but entirely missed in this study. For example, the putative adaptation of cranial morphology identified in NYC rats is likely polygenic [23]. Therefore, there is ample room for detecting targets and quantifying the mode of adaptation in urban populations in future research.

With the increasing urbanization of our planet, the effects of human activity on animals that inhabit cities merit further attention. Genetic adaptation and phenotypic plasticity have both been suggested as crucial drivers of success for rats and other species in urban settings—including great success in their capacity as human pests. Our results suggest that in the case of NYC brown rats, this adaptation—at least in part—could be rooted in rapid changes in the genetic composition of the population through natural selection.

## Acknowledgements

This work was funded by a fellowship from the Simons Society of Fellows (#633313) and a fellowship from the Stanford Center for Evolutionary and Human Genomics (CEHG) to A.H., National Science Foundation (NSF) grant DBI-1458059 to N.A.R. and D.A.P. and NSF grants DEB 1457523 and MRI 1531639 to J.M.-S. We thank Emily Puckett for her assistance in implementing demographic models from [19]. We thank Andrés Bendesky, Zach Fuller, Ana Pinharanda, Molly Przeworski and Nasa Sinnott-Armstrong for comments on the manuscript.

## Methods

### Data Availability

A Variant Calling Format (VCF) file that includes both the NYC sample and the Chinese sample, as well as all Supplementary Files can be found at https://doi.org/10.5061/dryad.08kprr4zn.

### Data collection

The NYC brown rat samples used in this study were a subset of nearly 400 rats collected previously for population genomic studies. Full details of this sampling effort are available in Combs et al. [20]. In brief, we trapped brown rats across the island of Manhattan in NYC between June 2014 and December 2015. Rats were trapped using lethal snap traps baited with a mixture of peanut butter, oats, and bacon, housed in bait stations (Bell Labs) and set for 24-hr periods. Traps were typically set outside earthen burrow systems or other areas with signs of ongoing rat activity. We then sampled 3-4 cm of tail tissue that was stored in 70% ethanol for downstream genetic analyses.

For comparison with the NYC sample, we used publicly-available whole genome sequences of 11 brown rats sampled from a 500 *km*^2^ area around the city of Harbin, Heilongjiang Province, China (European Nucleotide Archive ERP001276). Two BAM files were corrupt and our analysis is therefore based on 9 of the 11 Chinese samples. Heilongjiang Province is presumed to represent the ancestral range of *R. norvegicus*. This sample was previously used to examine the demographic history of brown rats [22]. Sequencing, mapping, variant calling and filtering were produced using a similar bioinformatic processing pipeline that we used for the NYC whole-genome samples and that is described below.

### Sequencing, read alignment, and variant calling

We sent RNAse-treated DNA extracts from 29 rat samples to the New York Genome Center for whole-genome resequencing on an Illumina HiSeq 2500. Rat samples were chosen to represent the diversity of rat colonies sampled across Manhattan (**Fig. 1**). The sequencing was designed to achieve at least 15X coverage for each sample. After sequencing, the NY Genome Center aligned cleaned and filtered reads to the *R. norvegivus* v. Rn 5.0 reference genome using BWA, and created a sorted, aligned BAM file using *Picard toolkit* [77]. *Duplicate reads were then removed using Picard toolkit*, followed by base quality score recalibration and local realignment around indels using *GATK* v3.7 [78].

We also used *GATK* for variant calling in both the NYC and Chinese samples. We made a file containing genotype information available for download from https://doi.org/10.5061/dryad.08kprr4zn in Variant Calling Format (VCF) for the *Rattus norvegicus* samples used in the study, including the 29 NYC individuals and the 9 Chinese individuals of the Deinum et al. study [22]. In the **Supplementary Materials**, we describe the bioinformatic procedures to produce this VCF.

### Selection statistics and calling candidate loci

For each SNP called in the NYC sample, we computed two statistics: G12 [33, 35], and H-scan [36]. For a sample of *n* diploids, both methods take as input *n* multi-site genotypes, where at each SNP, and for each individual, the two nucleotide alleles are replaced by pseudo-alleles corresponding to one of the (two or three) unique genotypes.

G12 examines a symmetric window of a fixed number of SNPs around each focal SNP. Denoting the frequency of the *k* unique multi-site genotypes at a window as *p*_1_, *p*_2_, *p*_3_, …, *p*_*k*_, ranked from most common to most rare (**Fig. 2A**), *G*_12_ is defined as

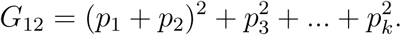

We used a window size of 201 SNPs to compute G12 values, conferring to a mean length of 29kbp and mean linkage disequilibrium (*r*^2^) of 0.135 between the SNPs at the ends of the window (**Fig. S10**).

H-scan is the mean pairwise identity tract length across all pairs of individuals in the sample. Namely, for a given focal site, let *h*_*ij*_ denote the length (in some distance metric, see below) of the maximal tract of pseudo-allele identity between individuals *i* and *j* that includes the focal site (and zero if no such tract exists—i.e. the pseudo-allele differs at the focal site). *H* is defined as the average of *h*_*ij*_ across all pairs,

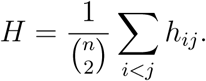

To run H-scan [36], a distance metric must be defined for the length of an identity-by-state tract for a given pair of individuals (*h*_*ij*_). We set the metric to be the number of SNPs rather than physical distance (base pairs) or genetic distance—an option that requires the input of a genetic map.

Lastly, we computed mean Weir-Cockerham per-SNP *F*_*st*_ in windows of 10kb with a 1kb step. In the **Supplementary Materials**, we provide further detail on pre-processing procedures, handling missing data and the method used to define candidate loci around top-scoring SNPs.

### Biological process enrichment analysis

In **Fig. 4** and **Fig. S3**, we examine whether top-scoring genes are enriched for any Gene Ontology (GO) biological process categories. We downloaded protein-coding gene annotations for the Rn5 reference genome from the UCSC genome browser [79]. We scored each gene by the maximal value of the statistic (G12 or H-scan) among focal SNPs inside the gene. We then ranked the genes by their score. **Table S4** (H-scan) and **Table S5** (G12) detail the gene scores, ranks, and associations. We considered a biological function category as enriched (considered as a “discovery”) if it was associated with at least three genes among the 50 top-ranking genes. In order to avoid spurious enrichment due to autocorrelated genotype homogeneity, we iteratively masked regions around genes in categories identified as enriched. Namely, with each new biological function category that we considered as enriched, we masked regions (i.e., excluded genes in these regions) that are within 30kb upstream and downstream of the three genes yielding the enrichment. This masking threshold is a modification of the masking approach we used for the genome-wide scans (**Supplementary Materials**), as we have found that without this modification, some gene clusters (e.g. the CYP2D cluster), that may also share biological process annotations, were over-represented due to shared G12 or H-scan signals.

We estimated the false discovery rate (FDR) for considering enriched categories among the top *r* ranked genes as “discoveries” (enriched biological processes). This FDR, appearing in grey in **Fig. 4** and **Fig. S3**, is the number of biological process categories that are expected to be associated with at least 3 of the genes ranked 1, …, *r* by chance alone, i.e. if ranks are independent of scores. Therefore, we simulated the null distribution of the number of discoveries by permuting scores across genes, reranking and counting the number of categories associated with at least 3 of the genes in the top *r* ranks. The FDR was then estimated as the average number of categories observed across 100 independent iterations of this permutation procedure. This permutation approach tests the null of independence of biological function and selection score, while controlling for the co-dependence of biological process categories by maintaining their joint distribution of the number of categories associated with each gene.

