## Supplementary Text and Figures for "Genetic Adaptation in New York City Rats"

### Contents

1 **Mapping and variant calling** 4

2 **Selection and diversity statistics** 5

3 **Relationship between genotype homogeneity and NYC-China differentiation** 7

4 **Calling candidate loci** 8

5 **H-scan in the Chinese sample at NYC candidates** 9

6 **Null distribution of distance from genes** 9

7 **Testing for enrichment of olfactory system genes** 10

8 **Neutral null simulations** 10

9 **Confirming the absence of known VKORC1 anticoagulant resistance-conferring alleles** 13


### List of Figures

S1 Properties of selection statistics. 14

S2 Demographic-model based null distribution of G12 15

S3 Biological processes enriched among top-scoring genes (with H-scan) 16

S4 Homogeneity in the Chinese sample scores at NYC-based candidates 17

S5 Selection statistics at VKORC1 18

S6 Genic polymorphisms in VKORC1 in NYC and Chinese rats. 19

S7 The distribution of G12 and H-scan. 20

S8 The distribution of G12 and H-scan across autosomes. 21

S9 Genome-wide H-scan vs. $F_{ST}$ scores . . . . . . . . . . . . . . . . . . . . . . . . . . 22
S10 Decay of linkage disequilibrium . . . . . . . . . . . . . . . . . . . . . . . . . . . . 23

### 1 Mapping and variant calling

We made three files containing genotype information available in the Variant Calling Format (VCF) for the rat samples used in the study: one for the 29 NYC rats, one for the 9 Chinese rats and a merged VCF containing both samples—38 individuals. Below, we describe the bioinformatic methods used to produce these VCFs.

We performed variant calling using GATK 3.7 [1] and following GATK's best practices [2, 3]. Before performing the variant calling, we edited the Rn5 reference genome file (downloaded from UCSC genome browser [4]) such that only contigs that had reads mapped to them were kept in the FASTA file. We then used GATK's genotypeGVCFs command to call variants as follows:

*java -Xmx80G -jar GenomeAnalysisTK.jar -T GenotypeGVCFs -R <fasta file> –variant <gCVF file> –variant <gCVF file> ... –variant <gCVF file> -o <raw-output.gvcf>*

This command was followed by subsequent commands to select only SNP variants and filter following GATK's best practices (as outlined in (`https://www.broadinstitute.org/partnerships/education/broade/best-practices-variant-calling-gatk-1`) as follows:

*java -jar GenomeAnalysisTK.jar -T SelectVariants -R <fasta file> -V <raw-output.gvcf> -selectType SNP -o <snp-output.vcf>*

*java -jar GenomeAnalysisTK.jar -T VariantFiltration -R <fasta file> -V <snp-output.vcf> –filterExpression "QD* $< 2.0$ | $FS > 60.0$ | *MQ* $< 40.0$ | *MQRankSum* $< -12.5$ | *ReadPosRankSum* $<$ *-8.0" −−filterName "best_prac_snp_filter" -o* $<$ *filtered-SNP.vcf* $>$

*java -jar GenomeAnalysisTK.jar -T SelectVariants -R <fasta file> -V <filtered-SNP.vcf> -env -o <analysis-ready.vcf>*

To generate a VCF file for the Chinese sample, we downloaded BAM files from the European Nucleotide Archive ERP001276 repository set up by Deinum et al. [5], we used *picard toolkit* commands sortSAM and reorderSAM and GATK's HaplotypeCaller followed by GenotypeGVCFs and filtering as for the NYC sample.

### 2 Selection and diversity statistics

For each SNP called in the NYC sample, we computed two statistics: G12 and H-scan. G12 [6] is computed exactly like H12 [7, 8], except that each individual is a diploid rather than a chromosome, and SNP genotypes (pairs of nucleotide alleles, e.g. C and T) are replaced by pseudo-alleles corresponding to unique genotypes, namely "A" for a homozygote C/C, "G" for a homozygote T/T and ". " for a heterozygote C/T. To compute G12, we first recoded the VCF of each autosome into a text file format accepted by a software designed to compute H12 downloaded from the Garud lab website (`https://garud.eeb.ucla.edu/selection-scan-scripts/`) [9]. Each row corresponds to a single SNP, and is written in the following comma-seperated fashion:

*<chromosomal position>,<allele of individual 1>,...,<allele of individual n>*

where the alleles in each row are 2 or 3 pseudo-alleles in our case, coded arbitrarily as "A","G" or ".", and corresponding to the unique genotypes at a given SNPs [9]. We filtered out sites that are invariant among the 29 rats (these may still appear in the VCF when all of the individuals were called as being homozygote for the non-reference allele). We also filtered sites where 7 individuals or more have unknown genotypes ("N"; these are predominantly low-coverage sites). When determining whether two multi-site genotypes of two individuals are identical, G12 ignores SNPs containing missing data for at least one of the individuals. We computed G12 by running H12 on the pseudo-allele text file ("genotype file") of each autosome $i \in \{1, ..., 20\}$ using the following command:

*python H12_H2H1.py <genotype file for chr i> 29 -o <G12 chr i> -w 200 -d 0*

where we specified that the score should be computed for all SNPs in the file and computed using windows of 201 SNPs.

To run H-scan we downloaded and compiled the C code on the Messer lab website [10]. We followed the practice suggested in the README file and ran the following command line for the genotype file of each chromosome $i$:

*H-scan -i <genotype text file> -d 1 > <H-scan result chr i>*

This command specifies that the score will be computed for all SNPs in the file, and uses the default setting of H-scan of measuring pairwise haplotype sharing using the number of SNPs rather than physical (basepair) or genetic distance (cM; an option that requires the input of a genetic map). Following our coding of missing data as dots ("."), H-scan treated missing data as unique pseudo-alleles and terminated identity tracts at SNPs where missing data was observed in at least one of the two individuals considered.

We computed the mean Weir-Cockerham $F_{st}$ across SNPs in windows of 10kb between the Chinese and NYC sample with the following *VCFTOOLS* v. 0.1.13 [11] command:

*vcftools –vcf <Joint NYC and Chinese samples VCF FILE> –weir-fst-pop <text file with NYC individuals identifiers> –weir-fst-pop <text file with chinese individuals identifiers> –fst-window-size 10000 –fst-window-step 1000 –out <Fst-scores.txt>*

**File S1** contains figures showing the values of the three statistics in each chromosome, and **Fig. S8** shows the distributions of G12 and H-scan values at each chromosome.

We estimated genome-wide nucleotide diversity ($\pi$) in the NYC and Chinese samples separately. We estimated $\pi$ as the average of estimates across 10kb windows in all autosomes, using the following *VCFTOOLS* v. 0.1.13 [11] command:

*vcftools –vcf <sample VCF FILE> –window-pi 10000 –out <pi-output>*

We estimated the standard errors for nucleotide diversity using a leave-one-out jackknife procedure.

Lastly, we estimated linkage disequilibrium (measured as the squared correlation in allelic states) between pairs of SNPs to inform our choice of the fixed analysis window size for G12. For computational efficiency, we only considered a random subset of $0.5\%$ of SNP pairs at a distance smaller than 1Mbp from each other. The genomewide trend is shown in **Fig. S10**. We used the following command in *plink* v. 1.9 [12]:

*plink –vcf <sample VCF FILE> –thin* $.005$ *–r2 –ld-window-kb 1000 –ld-window-r2 0 –ld-window 99999 –out <LD-output>*

### 3 Relationship between genotype homogeneity and NYC-China differentiation

In addition to the signal of high multi-site genotype homogeneity (measured with H-scan and G12), we have also identified regions with high genetic differentiation between the NYC sample and the Chinese sample as candidate targets of selection. We measured genetic differentiation using $F_{st}$ [13]. Genome-wide, $F_{st}$ is correlated with genotype homozygosity (Spearman $\rho = -0.46$, $p < 2 \times 10^{-22}$ for $F_{st}$ and H-scan averaged in windows of 10kb; **Fig. S9**). This correlation could likely be explained through the positive relationship between nucleotide homozygosity and the upper bound on attainable values of $F_{st}$ [14, 15].

To aid in the interpretation of candidates identified with each of the scans, we examined the values of $F_{st}$ in genotype homogeneity candidates and vice-versa. If genotype homogeneity candidates mostly correspond to selective sweeps that have occurred after the split from the ancestral population, then we expect high $F_{st}$ values in these loci. Indeed, $F_{st}$ values are elevated (80th percentile for H-scan, 76th percentile for G12) in genotype homogeneity candidates, consistent with a selective sweep elevating local differentiation from the Chinese population. However, the relationship between nucleotide homozygosity and $F_{st}$ implies that, even in the absence of selective sweeps, we should expect elevated $F_{st}$ values at NYC multi-site homogeneity peaks, and therefore the comparison to the genome-wide baseline may not be appropriate.

Conversely, genotype homogeneity values at $F_{st}$ candidates—70th percentile for H-scan but only 36th percentile for G12—might suggest that fewer of the $F_{st}$ candidates have undergone recent selective sweeps (namely, within the time-frame G12 and H-scan, applied to our sample size, are powered to detect sweeps in). $F_{st}$ candidates might still reflect older selective events (which no longer show strong haplotype structure), but they could also reflect a locally elevated mutation rate (**Fig. S1A**).

### 4 Calling candidate loci

For each one of the three statistics (G12, H-scan, $F_{st}$) we called the top 100 genomic intervals as "top-scoring loci". These loci are windows around top-scoring SNPs where the score (statistic) is high compared to the rest of the genome. We wished to avoid identifying multiple high-scoring SNPs that are in linkage, as they might represent the same adaptive event. We therefore used the following greedy clumping algorithm:

0. Set list of selectable SNPs to all SNPs on the chromosome
1. For i in 1...100
   1.1 Define the locus as the top scoring SNP among selectable SNPs.
   1.2 While SNP r, immediately downstream of the locus, has a score above threshold
      1.2.1 Add SNP r to locus
   1.3 While SNP l, immediately upstream of the locus, has a score above threshold
      1.3.1 Add SNP l to locus
   1.4 Remove set of locus SNPs—as well as all SNPs within $d$ base pairs upstream and $d$ base pairs downstream of the locus—from the set of selectable SNPs

The threshold used in Steps 1.2 and 1.3 was set to the top decile of scores in each chromosome.

The masking parameter $d$ (step $1.4$) was was set to 100kb in the analysis presented in the main text (including **Tables S1-3**). We performed a sensitivity analysis with $d$ set to 30kb instead, and find that it has a very minor effect on the results: all candidates called with $d = 100kb$ were also called with $d = 30kb$, in addition to two new loci with G12 and two new loci with H-scan (**Tables S7-9**).

### 5 H-scan in the Chinese sample at NYC candidates

In **Fig. 3D**, we show an example of a candidate locus found in our scan, based on the NYC sample. We show that in the same locus, in the Chinese sample, multi-site genotypes appear very heterogeneous. Here, we test whether a similar pattern is observed in other H-scan candidates. We computed H-scan in the Chinese sample (following the same procedures as those described for the NYC sample in **Section 2**). We then standardized the scores by substracting the means and dividing by the standard deviation in each chromosome separately. Genomewide, H-scan values calculated on the two samples independently are somewhat correlated (Pearson $r = 0.22, p < 10^{-16}$). In **Fig. S4** we examine the Chinese scores at NYC-based candidates. For most (12/17) of the candidates, the Chinese score is not significantly elevated (namely, an H score less than 2.5 standard deviations away from the chromosomal mean). This is consistent with the putative sweeps occurring in the ancestors of the NYC sample after the split from the ancestral population shared with the Chinese sample. Three of the remaining candidates—near the LOC680590, CHST11 and ST6GALNAC1 genes show elevated levels in the Chinese sample, but not as extreme as in NYC. The two remaining candidates—close to the XPR1 and GBGT1 genes—are more extreme outliers in the Chinese sample than in NYC.

### 6 Null distribution of distance from genes

In **Fig. 2C**, we compare the distance of candidate loci from genes in the rat genome to a null distribution of distances computed using 100 random loci in the genome. To generate a null distribution of 100 random loci, we performed a similar iterative masking procedure as in **Section S4** while maintaining the same locus size distribution. Namely, we used the following algorithm:

0. Set list of selectable SNPs to all SNPs in the genome

1. For i in 1...100

  1.1 Draw the position $x$ of a random SNP among selectable SNPs as the locus

  1.2 Let $d :=$ round(size of candidate locus $i$ in the real candidate loci list)/2.

  1.3 If $(x - d, x + d)$ only contains selectable SNPs, set $(x - d, x + d)$ as the locus else, go back to 1.1.

  1.4 Remove $(x - d, x + d)$ from the set of selectable SNP.

### 7 Testing for enrichment of olfactory system genes

For each of the three detection methods, we computed $x$, the number of candidate loci for which the nearest gene is an olfactory gene. We then calculated $y$, the number of null loci for which the closest gene is an olfactory gene. The 200 null loci were selected with the same algorithm as in **Section S6** after randomly permuting the selection statistic scores across SNPs. We then computed a p-value using Fisher's exact test on the following contingency table:

| | real data | null |
|---|---|---|
| olfactory loci | x | y |
| non-olfactory loci | 200-x | 200-y |

### 8 Neutral null simulations

To assess the significance of evidence for selection in our G12 scan, we initially used a formal hypothesis test with a simulation-based null distribution. We estimated the distribution of G12 under neutral dynamics, simulated under a demographic model recently proposed for NYC rats by Puckett et al. [16]. As we describe in the main text and below, we eventually decided not to use this approach to assess significance, but we include its description here for completeness.

We used the software *msms* to simulate the coalescent history of 58 chromosoms (29 pairs) [17, 18], using the demographic history for NYC rats inferred by Puckett et al. ("expansion" subpopulation in Figure 2 of [16]). The demographic model is shown in **Fig. S2A**. In our simulations, we specified a single site population mutation parameter of $\Theta = 4N_e\mu = 0.0017$—which we had estimated based on the rate of polymorphic sites ($K = 0.00684$) in our sample of 29 rats using Watterson's $\Theta$ estimator [19],

$$\Theta = \frac{K}{\sum_{i=1}^{28} i^{-1}}$$

We specified the population recombination parameter $= 4N_ec$ as $= 6.61$ where $c$ is the point recombination rate–following an estimated recombination rate of 0.6CM/Mb as estimated by Jensen-Seaman et al. [20], and $4N_e$ is estimated by $\Theta/\mu$ where $\mu = 9.29 \times 10^{-8}$ [16]. The length of the locus, $341,736bp$, was chosen to provide an expectation of 402 SNPs–twice the analysis window of 201 SNPs (empirically, all simulations indeed produced more than 201 SNPs). We used the middle SNP at the simulated locus in each simulation as the center of the analysis window. We used the following *msms* command to simulate 58 haploid rats with the parameters described above:

*java -Xmx500M -jar msms.jar 58 1 -t 102.365346125774 -r 6.61132483051282 -eN 0.0238129118689 0.350445827813644 -eN 0.056761725636 0.220879473122437 -eN 0.095724832725 0.185146576797008 -eN 0.133656902455 0.180591154299918 -eN 0.16683831618 0.192226504083746 -eN 0.194129281549 0.215752308177312 -eN 0.215888649953 0.250228986295763 -eN 0.233174026008 0.29606580322815 -eN 0.24726153957 0.354299697863844 -eN 0.2594189941 0.426234769965905 -eN 0.27089996417 0.513047304955648 -eN 0.283020340085 0.615238995235129 -eN 0.29723498528 0.732052176008236 -eN 0.31514948621 0.8610350128389 -eN 0.33850959688 0.998063828940034 -eN 0.36916842754 1.13798387142541 -eN 0.409016901645 1.27585943614117 -eN 0.45979121175 1.40866174920969 -eN 0.52263068523 1.53711653149769 -eN 0.59724692604 1.66744000399142 -eN 0.68070687135 1.81283393452849 -eN 0.7665512011 1.99408307560735 -eN 0.84639771867 2.23642739080683 -eN 0.91958162016 2.54849852826793 -eN 1.12735733248 2.57323577118922 -eN 1.3952916736 2.57323577118922 -eN 0.0065005514446 683.511037181462 -eN 0.0080437908448 683.511037181462*

*-eN 0.0099526157628 683.511037181462 > <output_file>*

We then randomly assorted the 58 haplotypes output by the simulation into 29 diploid individuals and computed $G12$ values.

We ran a total of 1,666,985 simulations to estimate the null distribution of G12, corresponding to five times the number of observed G12 data points. We report the neutral-simulation-based G12 values in **Table S6**. We computed a simulation-based p-value for all observed focal SNPs as the fraction of neutral simulations with a G12 value higher than the observed value. This approach gives a lower bound p-value of $\geq \frac{1}{1{,}666{,}985} = 6 \times 10^{-7}$, conservatively, for a candidate locus with a G12 value exceeding the maximal G12 values attained in simulations. After a Benjamini-Hochberg false discovery correction [21] combined with the conservative iterative-masking approach detailed in **Section 4**, all 20 G12 candidates that we report in **Table S1** are significant at $q < 2.4 \times 10^{-6}$.

We show a comparison of the genomewide distribution of G12 with the demography-based null in **Fig. S2B**. Taking the demography-based null model at face value, 0.07% of observed G12 values are higher than the maximal value attained across all simulations. Generally, the empirical distribution of G12 is highly dissimilar to the simulated one **Fig. S2B**. This discrepancy may be rooted either in a genome-wide deviation from the neutral model assumed in the *ms* simulations, a poor compatibility of the inferred demographic history to the complexities of the real population history, or a combination of the two. Of note, the demographic model may be particularly inaccurate in describing the recent history since the split from the ancestral population in China—which is likely an important factor driving the genome-wide distribution of statistics based on extended haplotype signatures.

Given the discrepancy between the genome-wide G12 distribution observed in the data and the simulations, we concluded that this model would be a poorly-calibrated null. We therefore take an alternative agnostic approach of focusing on the most extreme empirical values of our selection statistics as putative targets of adaptation. We nonetheless report the demography-based significance in **Table S1**.

#### 9 Confirming the absence of known VKORC1 anticoagulant resistance-conferring alleles

In the main text we discuss the absence of known resistance-conferring variants in our NYC sample. These include:

| Source | Variant |
|---|---|
| Rost et al. [22] | Ala21Thr<br>Ala26Thr<br>Arg33Pro<br>Arg35Pro<br>Tyr39Asn<br>Trp59Arg<br>Phe63Cys<br>Glu67Lys<br>Ile90Leu<br>Val112Leu<br>Leu128Gln<br>Tyr139Cys<br>Tyr139Phe<br>Tyr139Ser<br>Ile141Val<br>Ala143Val |
| Pelz et al. [23] | Leu120Gln<br>Ser56Pro |
| Iacucci et al. [24] | Ile123Ser |

All of these sites are not polymorphic in our sample. We have also confirmed that none of them appear in the Rn5.0 reference to which have mapped. Therefore, at all of the above sites, the non-resistant allele is fixed in our NYC sample.

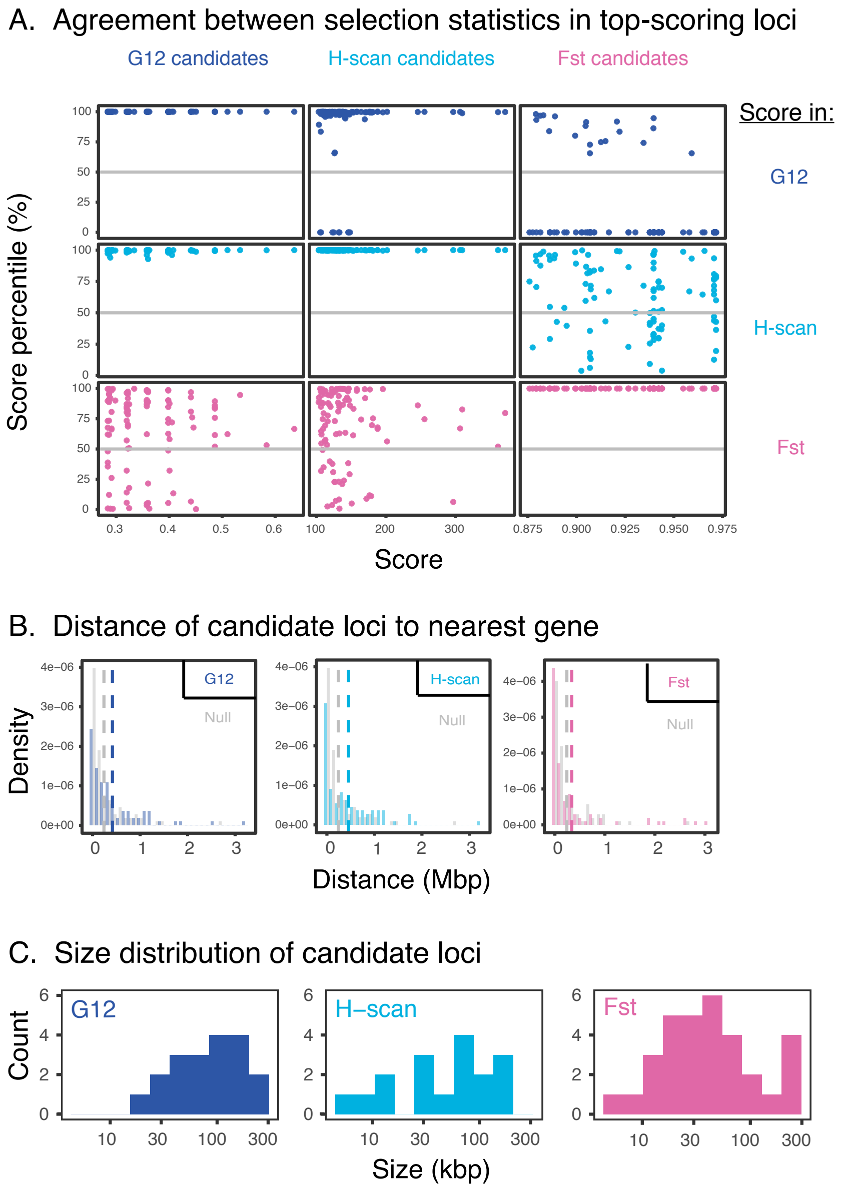


Figure S1: Properties of the selection statistics. **(A)** Chromosomal score percentile of each of the three statistics—in candidate loci identified with each of the three statistics. Circles show the 100 top-scoring loci for each statistic. G12 and H-scan agree on top chromosomal values. The correspondence between $F_{st}$ candidates and genotype homogeneity scores (and vice-versa) is weaker. **(B)** Distance from protein-coding genes. Color histograms show distances to the nearest gene for the 100 top-scoring loci of each statistic. Gray histograms show the null expectation, generated through a permutation of the candidate genes' locations. Vertical dashed lines show means. $F_{st}$, G12 and H-scan top-scoring loci are farther from genes than expected under the null, and $F_{st}$ top-scoring loci do not reject the null. **(C)** Size distribution of candidate loci. The boundaries of a candidate locus (around the focal test SNP) are set according to the algorithm in Section 4.

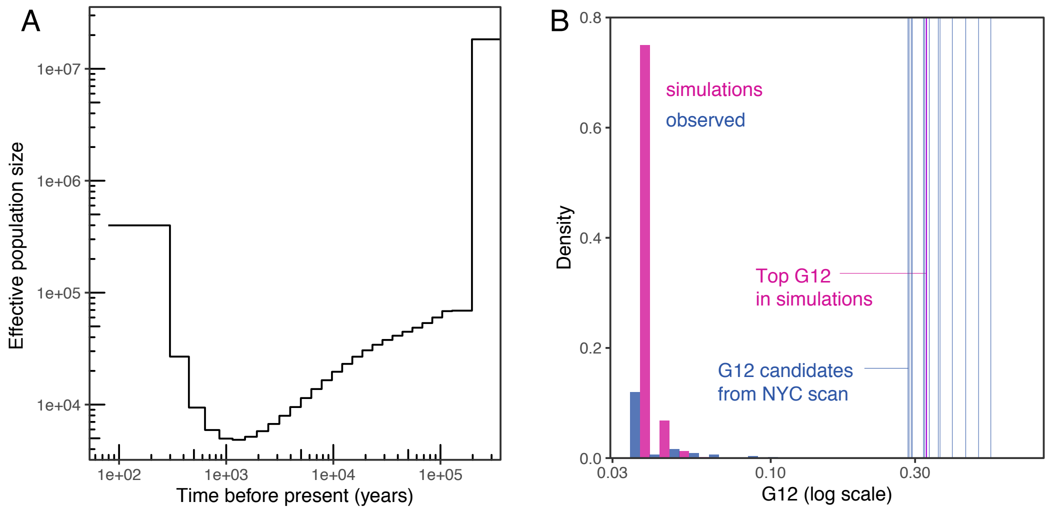


Figure S2: Demographic-model-based null distribution of G12. **(A)** A demographic model inferred by Puckett et al. [16], was used in simulations of the neutral evolution of NYC rat population. **(B)** A comparison of the distribution of G12 in neutral simulations (with the demographic model of panel A) and the observed data. The green vertical lines show the G12 values of all 20 G12-based candidates. The dark yellow vertical line shows the top G12 value in all 1,666,985 neutral simulation data points—five times the number of observed data points.

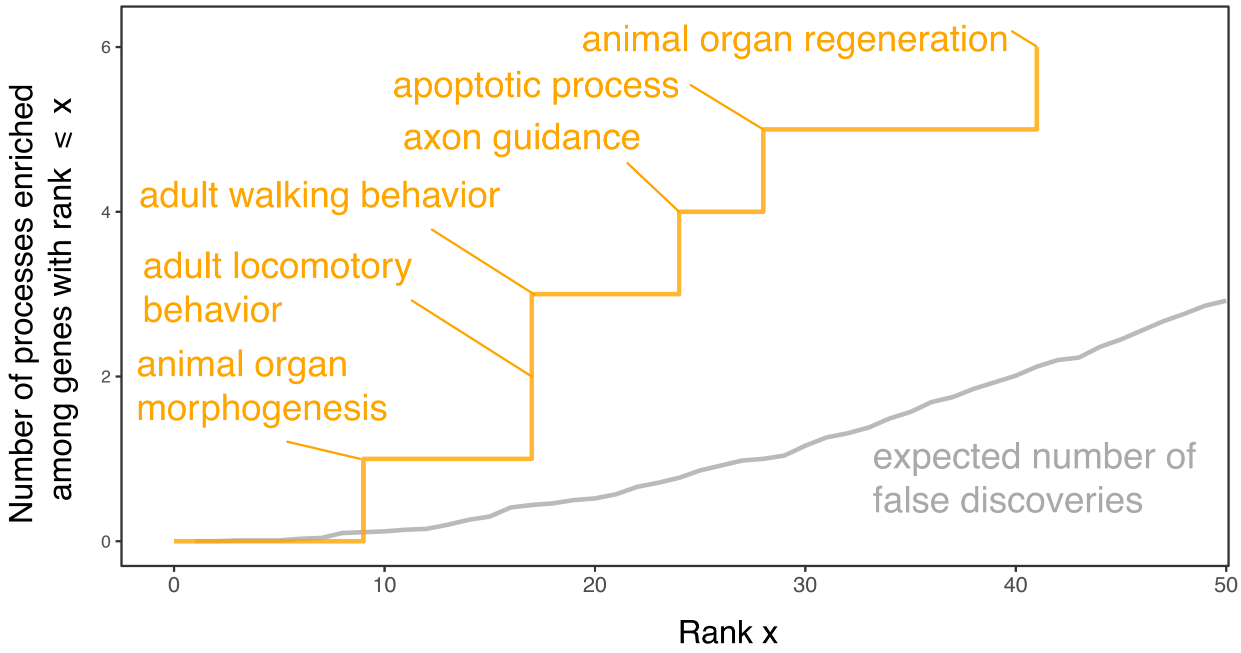


Figure S3: Biological processes enriched among top scoring genes (with H-scan). This figure mirrors Figure 4 of the main text, but is based on an analysis of H-scan rather than G12. We focus on candidate targets of adaptation arising from a comparison of H-scan values in genes. Each gene was given the maximum value of all focal SNPs in the gene, and genes were ranked by this maximal score. We consider Gene Ontology (GO) biological processes associated with at least three top-scoring genes as enriched, and the y-axis shows their cumulative count. Blue text labels show GO biological processes associated with exactly three of the (x value) top-scoring genes. The gray line shows the expected number of false (biological process) discoveries among the top X candidates, estimated based on permutations of gene scores.

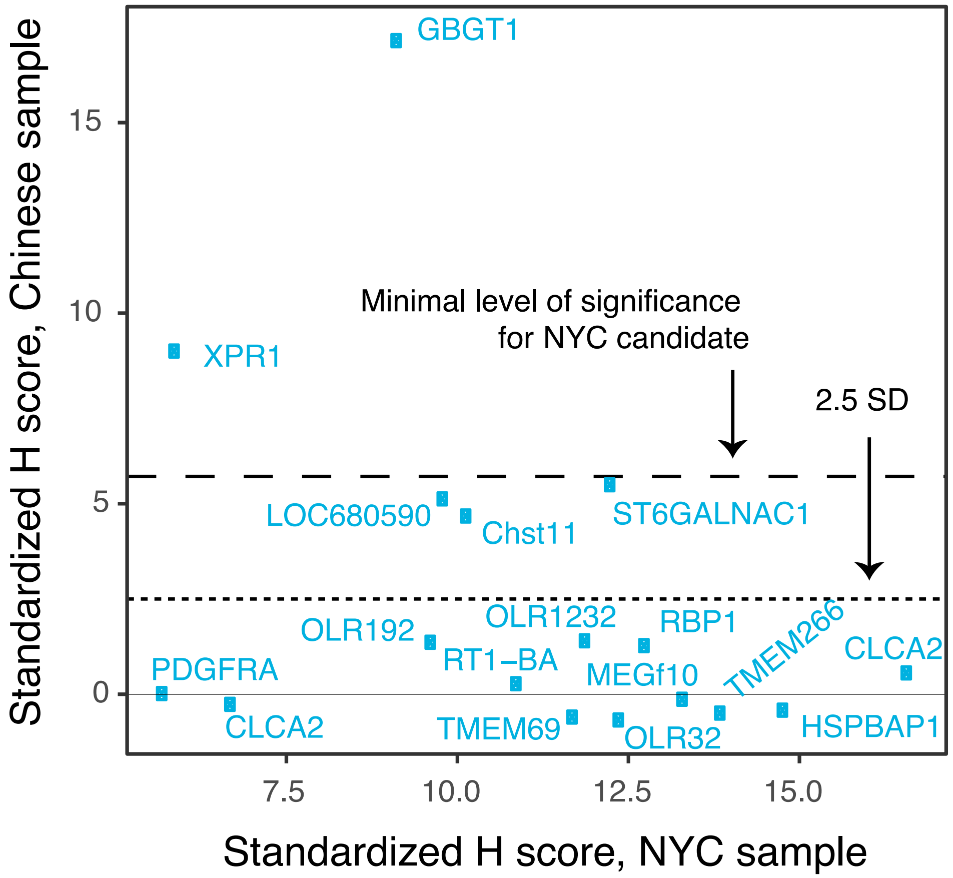


Figure S4: Homogeneity in the Chinese sample scores at NYC-based candidates. We computed H-scores (separately) on the sample of 9 rats from Harbin, Heilongjiang Province, China, and examined their values at H-scan candidates based on the scan in the NYC sample. H-scores (on both axes) are standardized for each chromosome separately, such that a value of 1 corresponds to an H score one standard deviation higher than the chromosomal mean. Labels near each data point show the name of the gene nearest to the candidate locus.

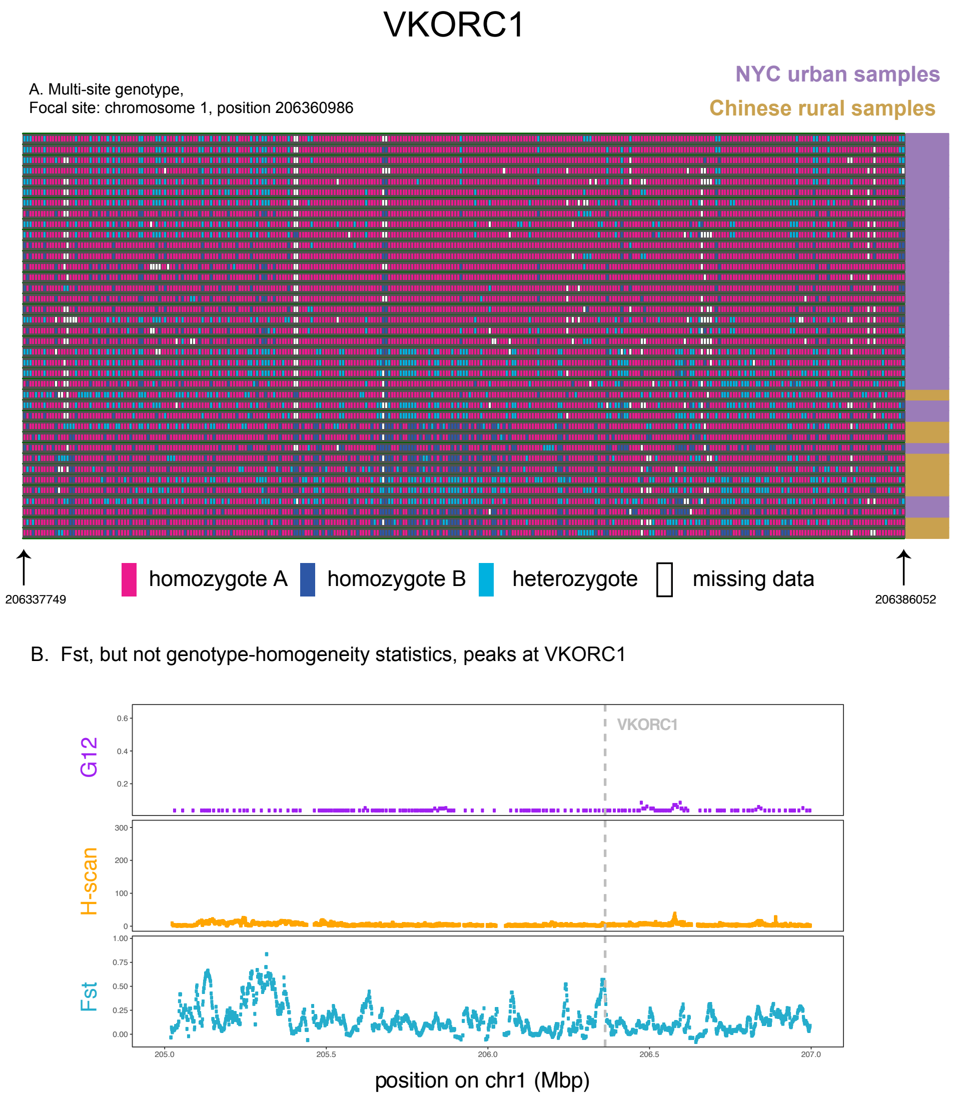


Figure S5: Selection statistics at VKORC1. **(A)** Multi-locus genotype composition around a SNP overlapping VKORC1, a gene in which mutations conferring Warfarin resistance have been suggested to have arisen through a selective sweep in different parts of the world. **(B)** $F_{st}$ reaches a high value (98th percentile in chromosome 1) at VKORC1. By contrast, H-scan and G12 values are not high near VKORC1.

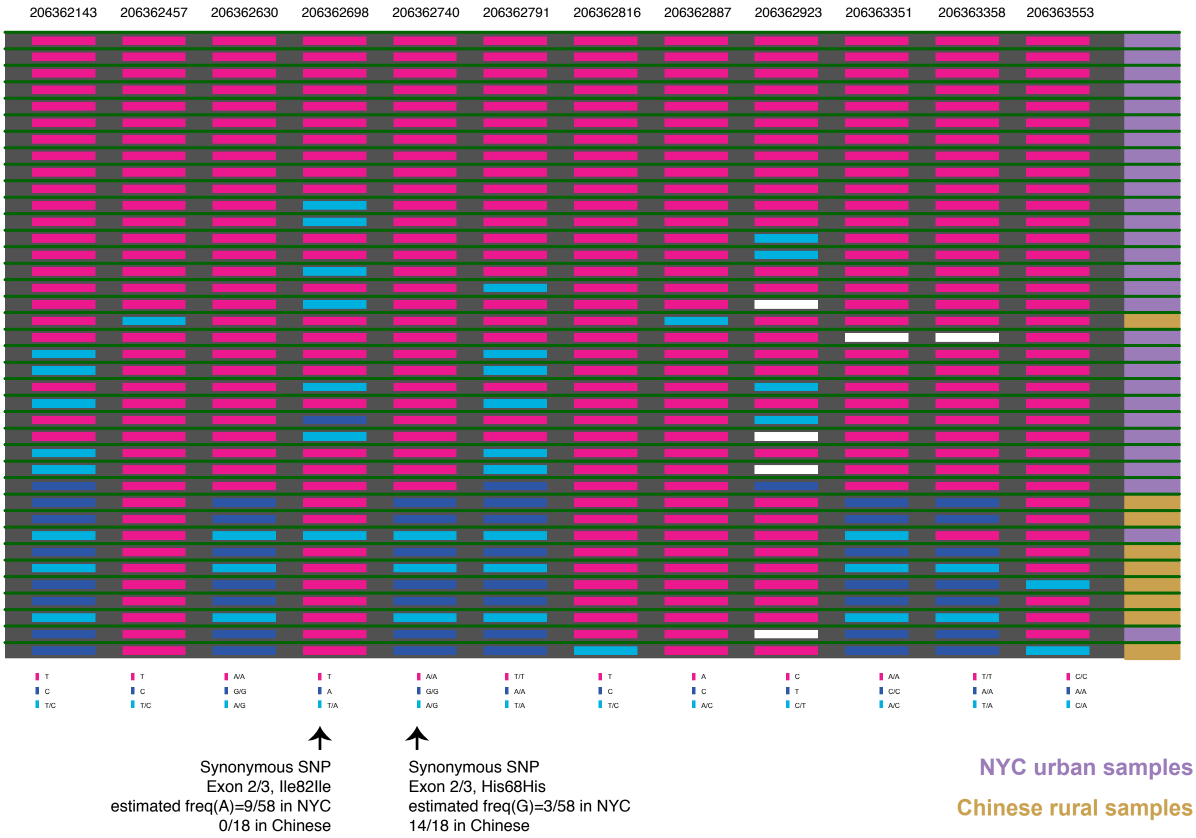


Figure S6: Genic polymorphisms in VKORC1 in NYC and Chinese rats. Shown are genotypes in all 12 SNPs identified in VKORC1: ten intronic and 2 synonymous SNPs. White rectangles denote missing data.

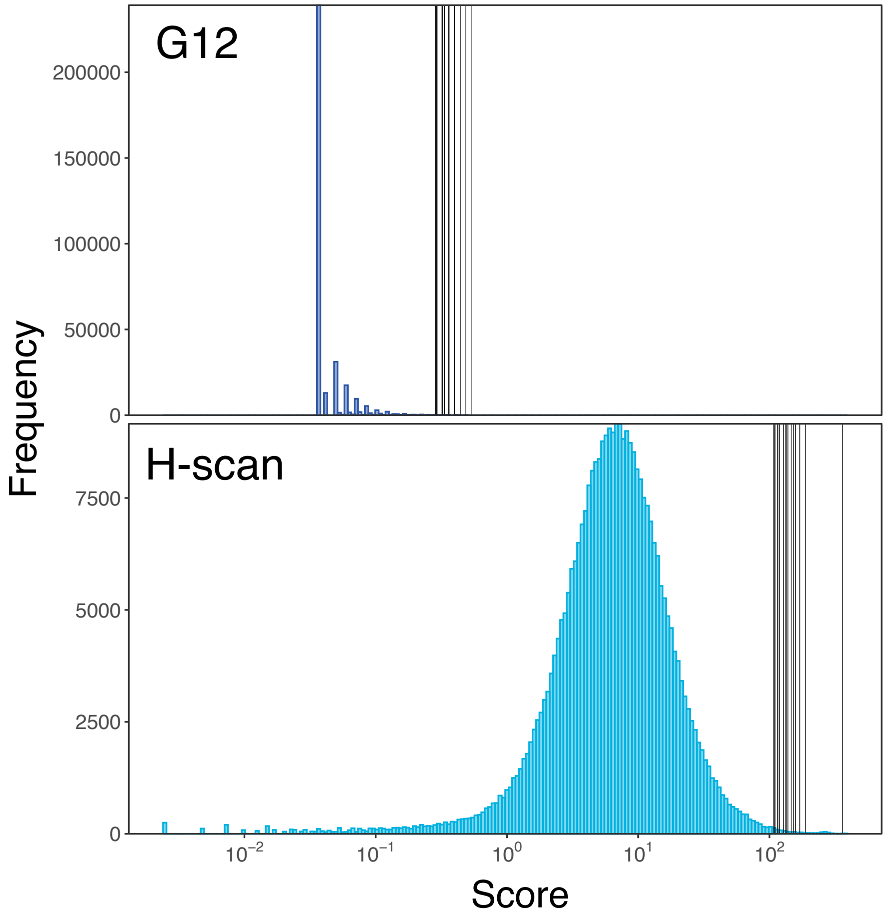


Figure S7: The distribution of G12 and H-scan. Black vertical lines show the values of candidates identified in each of the respective scans. Note that G12 values are from a limited set of values because they are functions of multi-locus genotype frequencies—sums of squares of between 1 and 28 positive numbers that sum to 1.

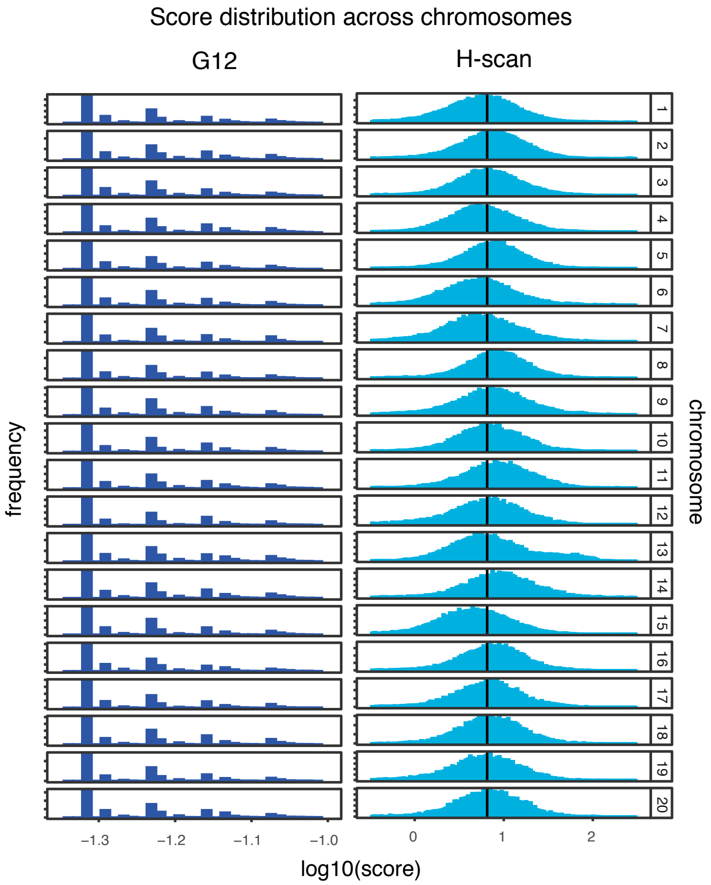


Figure S8: The distribution of G12 and H-scan across autosomes. Note that G12 values are from a limited set of values because they are functions of multi-locus genotype frequencies—sums of squares of between 1 and 28 positive numbers that sum to 1.

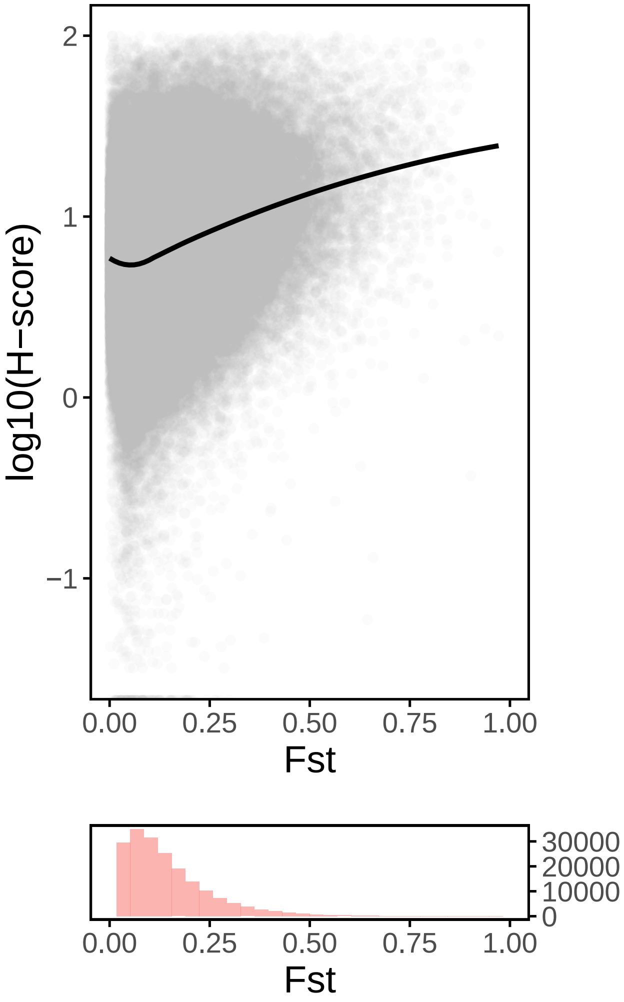


Figure S9: Genome-wide H-scan vs. $F_{ST}$ scores. The upper panel shows genome-wide H-scan vs. $F_{ST}$ values are shown in gray, and the black line shows a loess fit. $F_{ST}$ and H-scan are highly correlated (Spearman $\rho = -0.46, p < 2 \times 10^{-22}$ for averaged statistics in windows of 10kb). Both statistics are averaged within 10kb bins. The histogram in the bottom panel shows the marginal distribution of $F_{ST}$. Data points with estimated values of $F_{ST}$ below zero were excluded from this analysis.

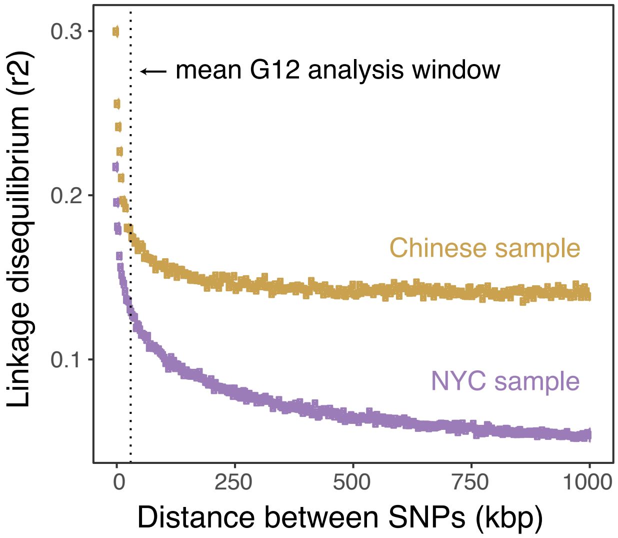


Figure S10: Decay of linkage disequilibrium. We estimated linkage disequilibrium–the squared correlation between the allelic states of pairs of sites–at distances ranging from 0-1Mbp apart. Each data point shows the average distance and linkage disequilibrium values of 10,000 pairs of sites, binned by distance. The dotted vertical line shows the mean analysis window size for G12, conferring to the fixed window size of 201 SNPs that we used in our selection scan.
